# *In vivo* antigen expression regulates CD4 T cell differentiation and vaccine efficacy against *Mycobacterium tuberculosis* infection

**DOI:** 10.1101/2021.02.02.429488

**Authors:** Helena Strand Clemmensen, Jean-Yves Dube, Fiona McIntosh, Ida Rosenkrands, Gregers Jungersen, Claus Aagaard, Peter Andersen, Marcel A. Behr, Rasmus Mortensen

**Author notes:** Author to whom correspondence should be addressed; Rasmus Mortensen, Department of Infectious Disease Immunology, Statens Serum Institut, Denmark.

## Abstract

New vaccines are urgently needed against *Mycobacterium tuberculosis* (Mtb), which kills more than 1.4 million people each year. CD4 T cell differentiation is a key determinant of protective immunity against Mtb, but it is not fully understood how host-pathogen interactions shape individual antigen-specific T cell populations and their protective capacity. Here, we investigated the immunodominant Mtb antigen, MPT70, which is upregulated in response to IFN-γ or nutrient/oxygen deprivation of *in vitro* infected macrophages. Using a murine aerosol infection model, we compared the *in vivo* expression kinetics of MPT70 to a constitutively expressed antigen, ESAT-6, and analysed their corresponding CD4 T cell phenotype and vaccine-protection. For wild-type Mtb, we found that *in vivo* expression of MPT70 was delayed compared to ESAT-6. This delayed expression was associated with induction of less differentiated MPT70-specific CD4 T cells but, compared to ESAT-6, also reduced protection after vaccination. In contrast, infection with an MPT70-overexpressing Mtb strain promoted highly differentiated KLRG1^+^CX3CR1^+^ CD4 T cells with limited lung-homing capacity. Importantly, this differentiated phenotype could be prevented by vaccination and, against the overexpressing strain, vaccination with MPT70 conferred similar protection as ESAT-6. Together our data indicate that high *in vivo* antigen expression drives T cells towards terminal differentiation and that targeted vaccination with adjuvanted protein can counteract this phenomenon by maintaining T cells in a protective less-differentiated state. These observations shed new light on host-pathogen interactions and provide guidance on how future Mtb vaccines can be designed to tip the immune-balance in favor of the host.

**Importance:** Tuberculosis, caused by Mtb, constitutes a global health crisis of massive proportions and the impact of the current COVID-19 pandemic is expected to cause a rise in tuberculosis-related deaths. Improved vaccines are therefore needed more than ever, but a lack of knowledge on protective immunity hampers their development. The present study shows that constitutively expressed antigens with high availability drive highly differentiated CD4 T cells with diminished protective capacity, which could be a survival strategy by Mtb to evade T cell immunity against key antigens. We demonstrate that immunisation with such antigens can counteract this phenomenon by maintaining antigen-specific T cells in a state of low differentiation. Future vaccine strategies should therefore explore combinations of multiple highly expressed antigens and we suggest that T cell differentiation could be used as a readily measurable parameter to identify these in both preclinical and clinical studies.

## Introduction

*Mycobacterium tuberculosis* (Mtb) has successfully survived in the human host and still causes 1.4 million deaths from tuberculosis (TB) disease annually (1). Formation of the granuloma is associated with the containment of infection as the environment within the granuloma suppresses Mtb growth in multiple ways, including oxygen and nutrient deprivation, exposure to acidic pH, and production of endogenous nitric oxide. In response to this, Mtb adapts by shifting between metabolic states often characterised by alterations in gene expression and thus changes in protein secretion and antigenic repertoire as well (2–5).

As part of the evolutionary adaptation, virulent strains across the mycobacterial tuberculosis complex (MTBC) vary in their expression of certain antigens, like the major secreted immunogenic protein 70 (MPT70) (6). Mtb produces very small amounts of MPT70 in *in vitro* cultures (7), but multiple studies have demonstrated that IFN-γ activation (5, 6) or starvation (8) induces MPT70 expression upon *in vitro* infection. Although MPT70 is a well-known immunodominant antigen during Mtb infection in both mice and humans (9–13), nothing is known about how MPT70’s *in vivo* antigen expression profile relates to the T cell phenotype it induces and its protective capacity as a vaccine antigen (13–16).

CD4 T cells are essential for protective immunity against Mtb (17–20) and there is mounting evidence that CD4 T cells develop into terminally differentiated IFN-γ producing effector T cells upon continuous antigen stimulation (21–23). The development of effector CD4 T cells is linked to sustained expression of markers and chemokine receptors associated with terminal Th1 differentiation and poor lung homing (24, 25). Less differentiated CXCR3^+^T-bet^dim^ CD4 T cells are able to enter the lung parenchyma and inhibit Mtb growth (25, 26) while terminally differentiated CD4 T cells co-expressing CX3CR1^+^KLRG1^+^ accumulate in the lung vasculature and provide no pulmonary control of Mtb infection (27, 28). Individual differences in antigen expression are suggested to shape T cell phenotype (22) and may therefore be a key determinant of vaccine protection.

The goal of this study was to investigate the impact of *in vivo* antigen expression on antigen recognition kinetics and adaptive immunity during Mtb infection, with MPT70 as a unique tool. Using the well-described 6kDa early secretory antigenic target (ESAT-6) as prototypic immunodominant model antigen (22, 29–31), we show that MPT70 displays delayed *in vivo* antigen expression as well as delayed immune recognition. This is associated with the induction of less differentiated CD4 T cells, but also lower protection in mice vaccinated with MPT70. Based on these observations, we hypothesise that high constitutive antigen expression is associated with increased T cell differentiation, but also improved vaccine capacity. In support of this, we demonstrate that artificial overexpression of MPT70 leads to accelerated CD4 T cell differentiation and diminished lung-homing capacity. However, vaccination with MPT70 counteracts this by stabilising a low degree of T cell differentiation and increases protection substantially in the MPT70 overexpressing strain compared to wild-type (WT) Mtb. Our study therefore reveals that antigen expression kinetics regulates CD4 T cell differentiation during infection and establishes a link between *in vivo* antigen expression, T cell differentiation, and vaccine protective capacity. This has implications for rational vaccine design, and future efforts in TB antigen discovery might use antigen-specific T cell differentiation as a readily measurable proxy for high *in vivo* antigen expression and increased vaccine potential.

## Results

### Delayed *in vivo* antigen transcription results in late immune recognition of MPT70

Previous studies indicate that MPT70 expression by Mtb is very low during *in vitro* cultivation (7, 11) but that expression is induced upon IFN-γ activation (5, 6) or nutrient-deprivation (8). Based on these studies we hypothesised that *in vivo* transcription and immune recognition of MPT70 would be delayed compared to ESAT-6, a constitutively expressed virulence factor (32, 33).

To map the kinetics of MPT70 expression *in vivo*, we infected a group of CB6F1 mice with Mtb Erdman, which we expected to produce low amounts of MPT70 (7). RNA was extracted from the post-caval lobe and cDNA was quantified by real-time qPCR using dual-labelled probes and normalised to 16srRNA. Expression levels of MPT70 and ESAT-6 mRNA were analysed prior to infection (week 0), at an early time point (week 4), and at a late time point (week 13). As expected, expression levels were below detection level prior to infection **(Figure 1a)**. At week 4, MPT70 expression was low and significantly lower than ESAT-6 but as the infection progressed to week 13, MPT70 expression increased and approached levels of ESAT-6, indicating a delayed expression profile **(Figure 1a)**.

**Figure 1.**
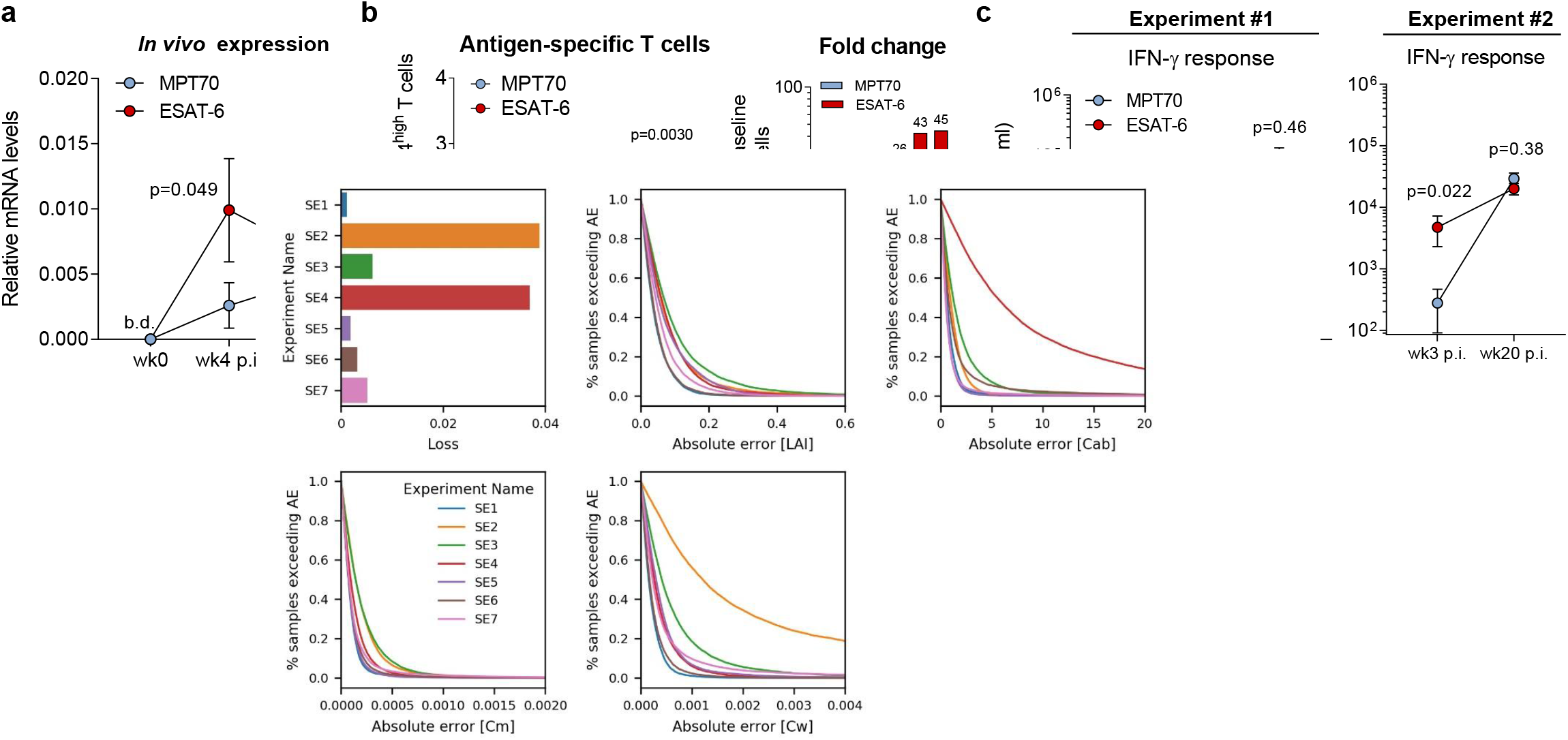
*In vivo* antigen expression and immune recognition of MPT70 is delayed during Mtb Erdman infection. CB6F1 mice were infected by the aerosol route with Mtb Erdman. **(a)** MPT70 and ESAT-6 *in vivo* gene expression were assessed pre-infection (week 0) and 4 and 13 weeks post-infection (p.i.) (n=4). The expression pre-infection was below detection levels (b.d.). Shown as average mean ± SEM. Paired t-test, two-tailed. **(b, left)** At week 3, 12, and 20 post Mtb infection, lungs were harvested for immunological analyses. Frequency of cytokine-producing CD3^+^CD4^+^ T cells specific for MPT70, ESAT-6 for the same time points as in c, exp1 (medium cytokine production subtracted) analysed by flow cytometry using **antibody panel 2** (n=4). Shown as average mean ± SEM. One-way ANO-VA with Tukey’s Multiple Comparison test **(b, right)** Fold change in cytokine-producing CD4 T cells from baseline. **(c)** Lung cells from infected mice were restimulated *in vitro* with media, MPT70 or ESAT-6 for five days. Culture supernatant was harvest and measured for IFN-γ levels in two individual experiments (n=4). Values were log-transformed and shown as average mean ± SEM. Paired t-test, two-tailed.

We next investigated the kinetics of the immune recognition to the two antigens during the course of infection. Mice were infected as previously described and antigen-specific immune responses were detected at 3, 12, and 20 weeks post-infection, either by intracellular cytokine staining (ICS) measuring the frequency of antigen-specific CD4 T cells producing IL-2, TNF-α, or IFN-γ **(Figure 1b and Figure S1)** or IFN-γ release in cultures of stimulated splenocytes **(Figure 1c)**. Notably, in two independent experiments, we observed that the immune recognition of MPT70 was very low in the early phase of infection but continued to increase as the infection progressed to week 12 and 20 **(Figure 1b and Figure 1c)**. This was in contrast to ESAT-6 responses that were greater at week 3 and 12, after which they plateaued.

Together, these data indicate that *in vivo* expression of MPT70 is delayed compared to ESAT-6 and that this difference in kinetics is associated with delayed onset of specific CD4 T cell responses.

### MPT70-specific CD4 T cells maintain a low degree of differentiation

Continuous stimulation with high levels of antigen is known to drive T cells towards terminal differentiation (21–23) and we therefore explored whether the delayed antigen availability of MPT70 favoured the development of less differentiated T cells during infection. In order to address this, we first characterised MPT70 and ESAT-6 specific T cells according to their expression of intracellular cytokines associated with Th1 differentiation. As previously defined (22, 34), a functional differentiation score (FDS) represents a simple measure for a T cell’s differentiation status and is calculated as the ratio of all highly differentiated IFN-γ producing T cell subsets divided by less differentiated T cell subsets producing other cytokines (IL-2, TNF-α). An FDS score >1 is therefore indicative of a response with more highly differentiated T cells than less differentiated T cells. During the first two weeks of infection, MPT70 and ESAT-6 specific CD4 T cells displayed similar FDS in the range of 2. From week two to four, the FDS of both T cell subsets increased to 3.8 and 5.9, respectively. From week 4 and onwards, the FDS of ESAT-6 T cells continuously increased to reach 18, while the FDS of MPT70 T cells remained constant around 4, denoting that MPT70 CD4 T cells are not driven towards terminal differentiation to the same extent as ESAT-6 **(Figure 2a)**. In TB infected mice, CXCR3^+^KLRG1^-^Tbet^dim^ T cells migrate into the lung parenchyma and control the infection (26, 28), while intravascular (iv) T cells have a high expression of KLRG1, CX3CR1, and T-bet (24). In accordance with the FDS data, we observed a substantially lower proportion of cytokine expressing KLRG1^+^ CD4 T cells after MPT70 stimulation compared to ESAT-6, and this difference was sustained throughout the entire infection **(Figure 2b)**. Investigating the ability of these CD4 T cell subsets to enter the infected lung tissue by CD45 iv staining further supported that a smaller fraction of MPT70-specific CD4 T cells were retained in the lung-associated vasculature (CD45 iv^+^) compared to ESAT-6 CD4 T cells **(Figure 2c)**.

**Figure 2.**
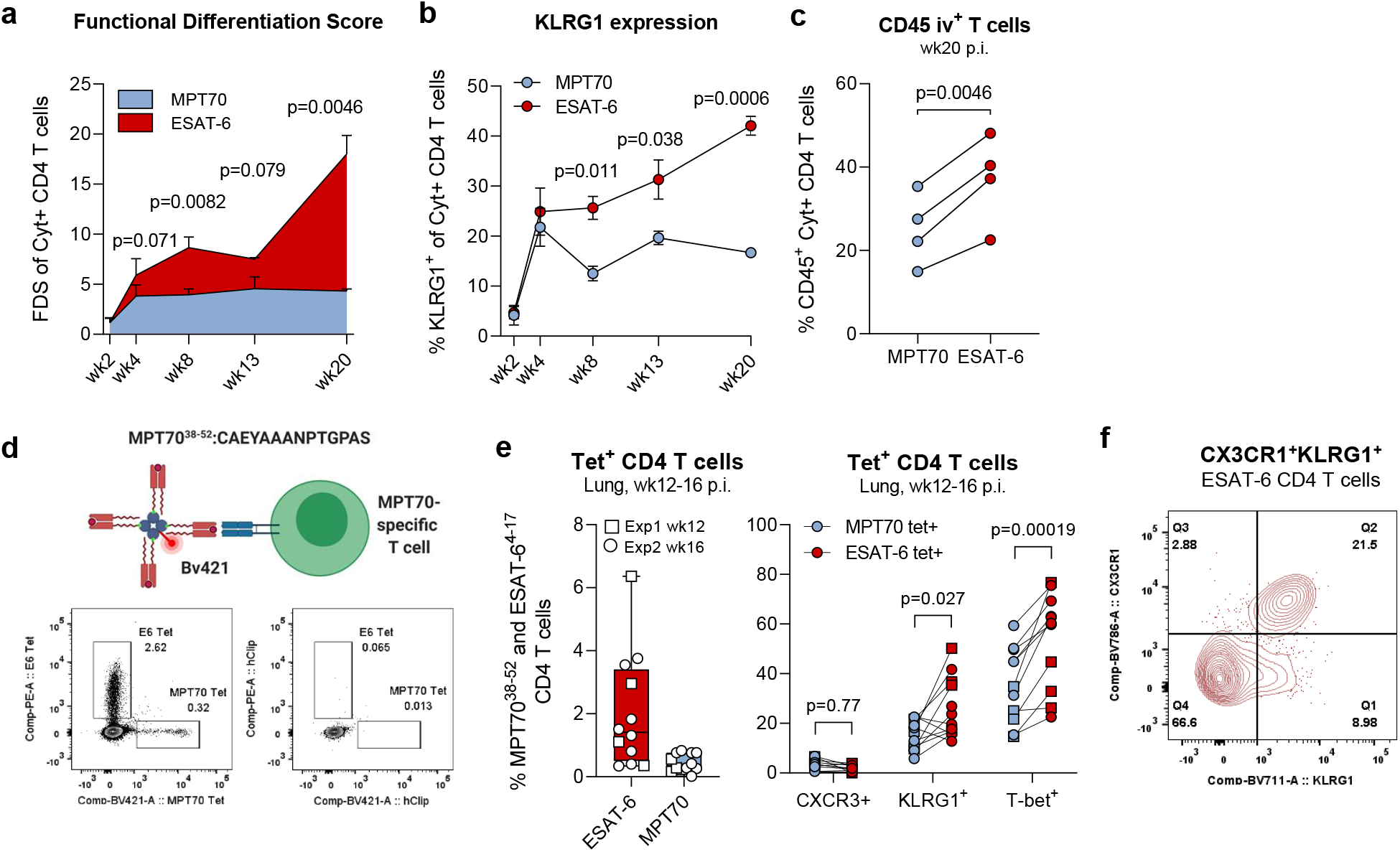
MPT70-specific CD4 T cells maintain a low differentiation state compared to ESAT-6. **(a)** The functional differentiation score (FDS) of MPT70 and ESAT-6-specific CD4 T cells over the course of Mtb Erdman infection (n=4). The FDS is defined as the ratio of all IFN-γ producing CD4 T cell subsets divided by subsets producing other cytokines (IL-2, TNF-α), but not IFN-γ (high FDS = high IFN-γ production). Multiple t-tests with correction for multiple testing using the Holm-Sidak method. Shown as average mean ± SEM. Flow Cytometry gating as depicted in **Figure S1**, using **antibody panel 2**. **(b)** Frequencies of KLRG1 expressing MPT70 and ESAT-6-specific CD4 T cells throughout infection (n=4). Shown as average mean ± SEM. Multiple t-tests with correction for multiple testing using the Holm-Sidak method. **(c)** Frequency of CD45-labelled MPT70 and ESAT-6 specific CD4 T cells in the lung-associated vasculature (CD45^+^) 20 weeks post-infection (p.i.) with Mtb (n=4). Paired t-test, two-tailed**. (d, upper)** Schematic representation of custom-made I-A^b^:MPT70_38-52_ MHC-II tetramer. **(d, lower)** Representative concatenated FACS plots showing frequencies of I-A^b^:MPT7 0_38-52_ and I-Ab:ESAT-6_4-17_ tetramer^+^ CD4 T cells or corresponding hClip tetramer^+^ CD4 T cells in lungs of mice 12 weeks post Mtb infection (n=4). **(e)** Frequency of I-A^b^:MPT7 0_38-52_ and I-Ab:ESAT-6^4-17^ CD4 T cells 12-16 weeks post Mtb infection expressing CXCR3, KLRG1, and T-bet. Parametric, paired t-test, two-tailed (n=12). Flow Cytometry gating as depicted in **Figure S3** using **antibody panel 1. (f)** Concatenated FACS plot of CX3CR1^+^KLRG1^+^ co-expressing ESAT-6^4-17^ CD4 T cells (n=4).

We next wanted to confirm these observations using an MHC-II tetramer. In contrast to ICS, tetramers identify antigen-specific T cells without the risk of affecting the expression of certain markers due to *ex vivo* stimulation. We therefore epitope mapped the MPT70 protein (29) and developed a murine MHC-II tetramer specific for I-A^b^:MPT7 0_38-52_ **(see method section, Figure 2d, Figure S2)**. In the lungs of mice infected with Mtb for 12-16 weeks, we found an average of 2.02% tetramer-positive I-A^b^:ESAT-6_4-17_ and 0.44% I-A^b^:MPT70 _38-52_ specific CD4 T cells **(Figure 2e)**. Exploring the expression of CXCR3, KLRG1, CX3CR1, and T-bet showed that MPT7 0_38-52_ specific CD4 T cells expressed significantly lower levels of KLRG1 (p=0.027) and T-bet (p=0.00019) compared to ESAT-6_4-17_ specific T cells **(Figure 2e)**. Although there was no difference in CXCR3 expression, the vast majority of KLRG1^+^ T cells coexpressed CX3CR1^+^, which is associated with vascular T cells (24, 25) **(Figure 2f)**, and therefore in agreement with the data obtained by CD45 iv staining.

In summary, these studies show, by both cytokine production pattern and expression of differentiation markers, that MPT70-specific CD4 T cells are maintained at a lower state of differentiation throughout infection compared to ESAT-6 induced T cells. This difference translates into a higher functional capacity of MPT70-specific CD4 T cells to migrate to infected lung tissue.

### The impact of vaccinating with MPT70 is lower than for ESAT-6

The previous data showed that T cells specific for MPT70 were less differentiated than those specific for ESAT-6 during experimental infection. We next investigated the significance of this in the context of vaccine efficacy. Mice were immunised three times with MPT70 or ESAT-6 in the cationic adjuvant formulation 1^®^ (CAF01^®^) adjuvant (35), and immune responses were characterised two weeks after the final vaccination. ICS of stimulated splenocytes detected 0.79 % MPT70-specific CD4 T cells compared to 0.42 % for ESAT-6, showing that both antigens were immunogenic and, if anything, MPT70 induced higher responses than ESAT-6 **(Figure 3a)**. However, 3 weeks after Mtb Erdman challenge there was a bigger proportion of ESAT-6-specific T cells in the lung compared to MPT70 T cells, indicating earlier expansion/recruitment of ESAT-6-specific T cells **(Figure 3b).** A characterisation of the MPT70 and ESAT-6 specific CD4 T cells showed that there was no difference in T cell differentiation pre-infection **(Figure 3c)**, indicating that there was no intrinsic antigen effect on this parameter. In contrast, a similar analysis post-infection revealed that vaccination with ESAT-6 had a greater impact on lowering the antigen-specific T cell differentiation in this setting **(Figure 3d)**. Similar to the observation in Figure 1a, MPT70-specific T cells had an FDS of around 2.7 during early infection and vaccination did not change this noticeably. In unvaccinated animals, this level was sustained at week 20, while vaccination with MPT70 lowered this to 1.1. In contrast, the FDS of ESAT-6 specific T cells remained high throughout infection (week 3 = 6.9 and week 20 = 5.8), while vaccination with ESAT-6 resulted in a substantially lower differentiation level around 0.8-1.8 throughout the infection **(Figure 3d)**. These observations were confirmed by KLRG1 staining, which showed a similar pattern to FDS **(Figure S4a)**. Finally, we determined the protective efficacy by plating lung homogenates. Of note, at weeks 3-4, we observed significant protection of both ESAT-6 (p<0.0001) and MPT70 (p=0.0001), demonstrating the vaccine potential of both antigens **(Figure 3e)**. However, over the course of four independent experiments, bacterial burdens were lower in ESAT-6 vaccinated animals than MPT70-vaccinated animals (p=0.047 at 3-4 weeks p.i.) **(Figure 3e and Figure S4b)**.

**Figure 3.**
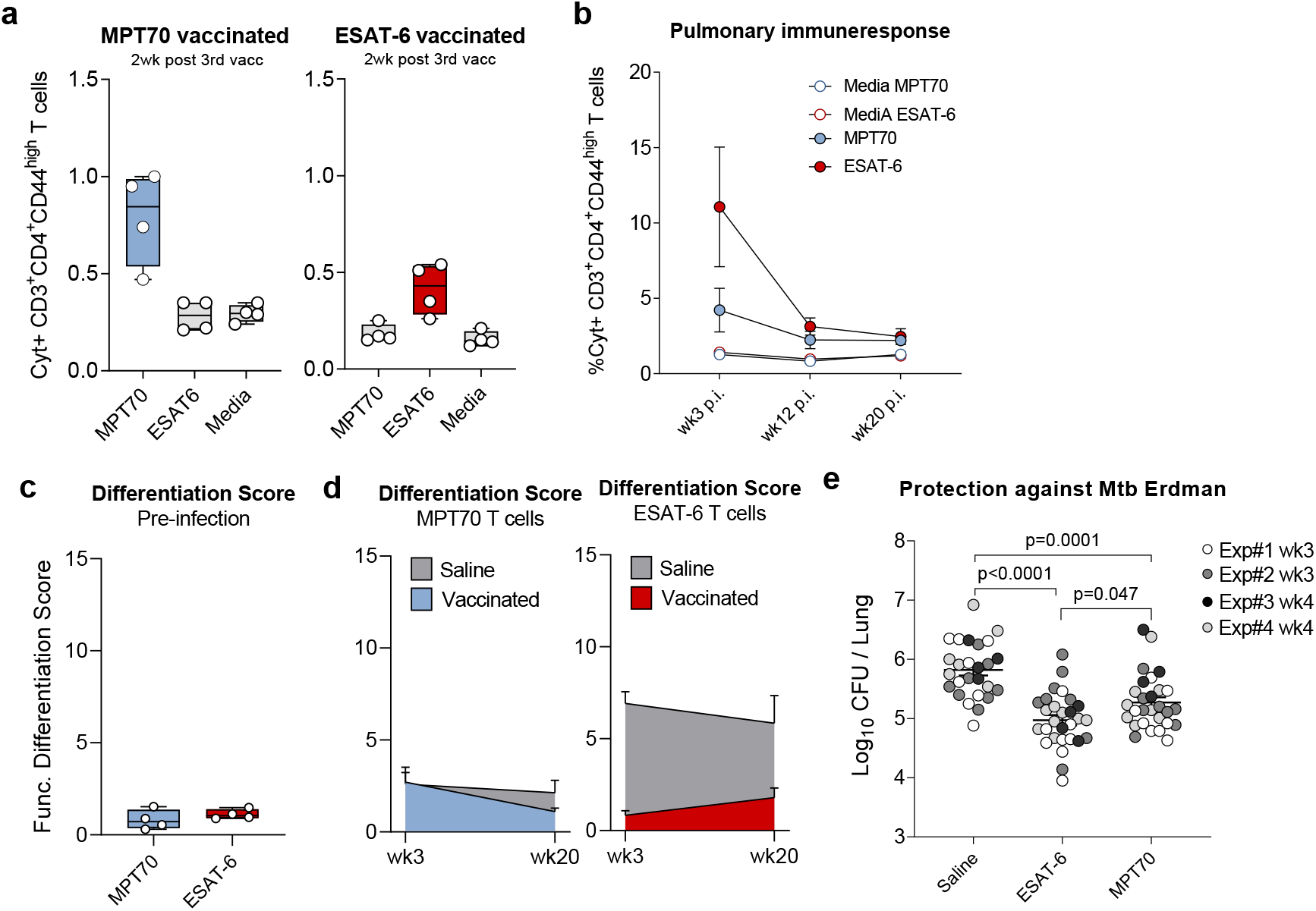
Vaccination with MPT70 has a lower impact on CD4 T cell differentiation than ESAT-6. Female CB6F1 mice were immunised with either MPT70 or ESAT-6 recombinant protein three times s.c. and challenged with Mtb Erdman six weeks post the third immunisation. **(a)** Frequency of MPT70 and ESAT-6 specific CD4 T cells in the spleen two weeks post the third vaccination (n=4). **(b)** Frequency of MPT70 and ESAT-6 specific CD4 T cells in the lung week 3, 12, and 20 post Mtb infection (n=4). Shown as average mean ± SEM. **(c)** Functional differentiation score (FDS) of MPT70 and ESAT-6-specific CD4 T cells pre-infection in the spleen (n=4). **(d)** FDS of MPT70 and ESAT-6-specific CD4 T cells 3 and 20 weeks post Mtb infection in lungs of vaccinated and saline mice (n=4). Shown as average mean ± SEM. Flow Cytometry gating as depicted in **Figure S1**, using **antibody panel 2. (e)** The bacterial burden was determined in the lungs of saline, MPT70 and ESAT-6 vaccinated mice at 3-4 weeks post Mtb infection (n=26-28). The graph represents four individual experiments of which experiment 4 is already published in (29). One-Way ANOVA with Tukey’s multiple comparison test.

Taken together, vaccination with ESAT-6 had the highest impact on T cell differentiation during subsequent infection and while vaccination with both antigens induced robust protection, there was better protection with ESAT-6. Although observed with antigens of different size and immunogenicity, this suggests that *in vivo* antigen expression could regulate T cell quality as well as protective capacity.

### Constitutive expression of MPT70 accelerate T cell differentiation and improve vaccine protection

To investigate whether T cell differentiation and vaccine protection is directly linked to *in vivo* antigen transcription, we utilised a recently engineered H37Rv strain (36) (herein called H37Rv::mpt70^high^) with significantly increased *in vitro* expression of MPT70 due to insertion of *sigK* (Rv0445c) and *rskA* (Rv0444c) from *M. orygis* (6, 36). In line with the previous report (36), we observed that this strain upregulated *in vitro* expression of MPT70 compared to wild type (WT) H37Rv, while very little changes were observed for the regulators of MPT70 transcription (SigK and RskA) **(Figure 4a)**. From this, we anticipated that the H37Rv::mpt70^high^ strain would have an increased early *in vivo* expression of MPT70. Transcription analysis of mRNA from lungs of mice 3 weeks after aerosol infection confirmed this, as MPT70 was 6.7 fold higher expressed by H37Rv::mpt70^high^ than WT H37Rv infected mice **(Figure 4b).** This analysis also confirmed the observations with Mtb Erdman in Figure 1a, showing that expression of MPT70 in WT H37Rv was very low at week 3 in contrast to ESAT-6 **(Figure S5a)**. Of note, complementation of H37Rv did seem to affect the bacterial fitness *in vitro* **(Figure S5b)**, which was also associated with a small, but detectable, difference in CFU at day 1 **(Figure 4c)**. Interestingly, H37Rv::mpt70^high^ and WT H37Rv had similar *in vivo* growth up until week 3, but overexpression of MPT70 seemed to impact long-term persistence negatively **(Figure 4c)**. We next analysed the impact on T cell responses in two independent experiments. Since bacterial load is expected to influence T cell differentiation (21–23), we focused our analysis around week 3 post-infection, where the number of bacteria was the same for WT H37Rv and H37Rv::mpt70^high^. The first experiment demonstrated that the CD4 T cell response against MPT70 was significantly increased in mice infected with H37Rv::mpt70^high^ compared to WT H37Rv (**Figure 4d)**. For comparison, ESAT-6 responses did not differ between the two strains **(Figure 4d)**. In agreement with this observation, a second experiment showed that the response to MPT70 vaccination was increased in mice infected with H37Rv::mpt70^high^ (p=0.0064) compared to WT infected mice **(Figure 4e)**. We then asked if the early expression and elevated MPT70 immune response accelerated CD4 T cell differentiation and altered expression of markers associated with lung homing. CXCR3 is primarily expressed on lung-homing T cells (25, 26), whereas CX3CR1 is associated with T cells in the vasculature (37). Studying these surface markers on MPT70-specific CD4 T cells revealed a substantially higher proportion of CX3CR1^+^KLRG1^+^ T cells after H37Rv::mpt70^high^ infection (20.6%) compared to H37Rv infection (11.8%) indicating increased differentiation and decreased lung homing capacity for H37Rv::mpt70^high^ primed MPT70 CD4 T cells **(Figure 4f)**. This was also evident in a vaccination setting, as MPT70 immunisation induced the biggest reduction of CX3CR1^+^KLRG1^+^ expressing T cells after H37Rv::mpt70^high^ infection **(Figure S5c)**. In contrast, the frequency of CX3CR1^+^KLRG1^+^ expressing ESAT-6 T cells was not different between the strains, demonstrating that the increased T cell differentiation was a specific effect of MPT70 overexpression **(Figure S5d)**. In line with increased MPT70-specific T cell differentiation, a lower frequency of CXCR3^+^ expressing CD4 T cells was found in H37Rv::mpt70^high^ infected animals (14.0%) compared to H37Rv infected (20.1%), which also correlated with a higher proportion of CD45-labelled MPT70 CD4 T cells located in the lung-associated vasculature (23.4% to 10.2%) **(Figure 4f)**. Together, the immune data showed that increased early *in vivo* expression altered the MPT70 specific immune responses to resemble the ones of ESAT-6 after WT Mtb infection. We finally addressed whether vaccine-induced protection of MPT70 was increased if mice were challenged with the H37Rv::mpt70^high^ strain. Immunised mice were challenged with either WT H37Rv or H37Rv::mpt70^high^ and bacterial numbers were determined in the lungs at weeks 3, 12, and 22. Consistent with data from Mtb Erdman **(Figure 3e)**, MPT70 vaccination conferred less protection than ESAT-6 against challenge with WT H37Rv **(Figure 4g)**. In contrast, MPT70 vaccination induced a substantial reduction in bacterial load at all time points in mice challenged with H37Rv::mpt70^high^, providing protection that was comparable to ESAT-6.

**Figure 4.**
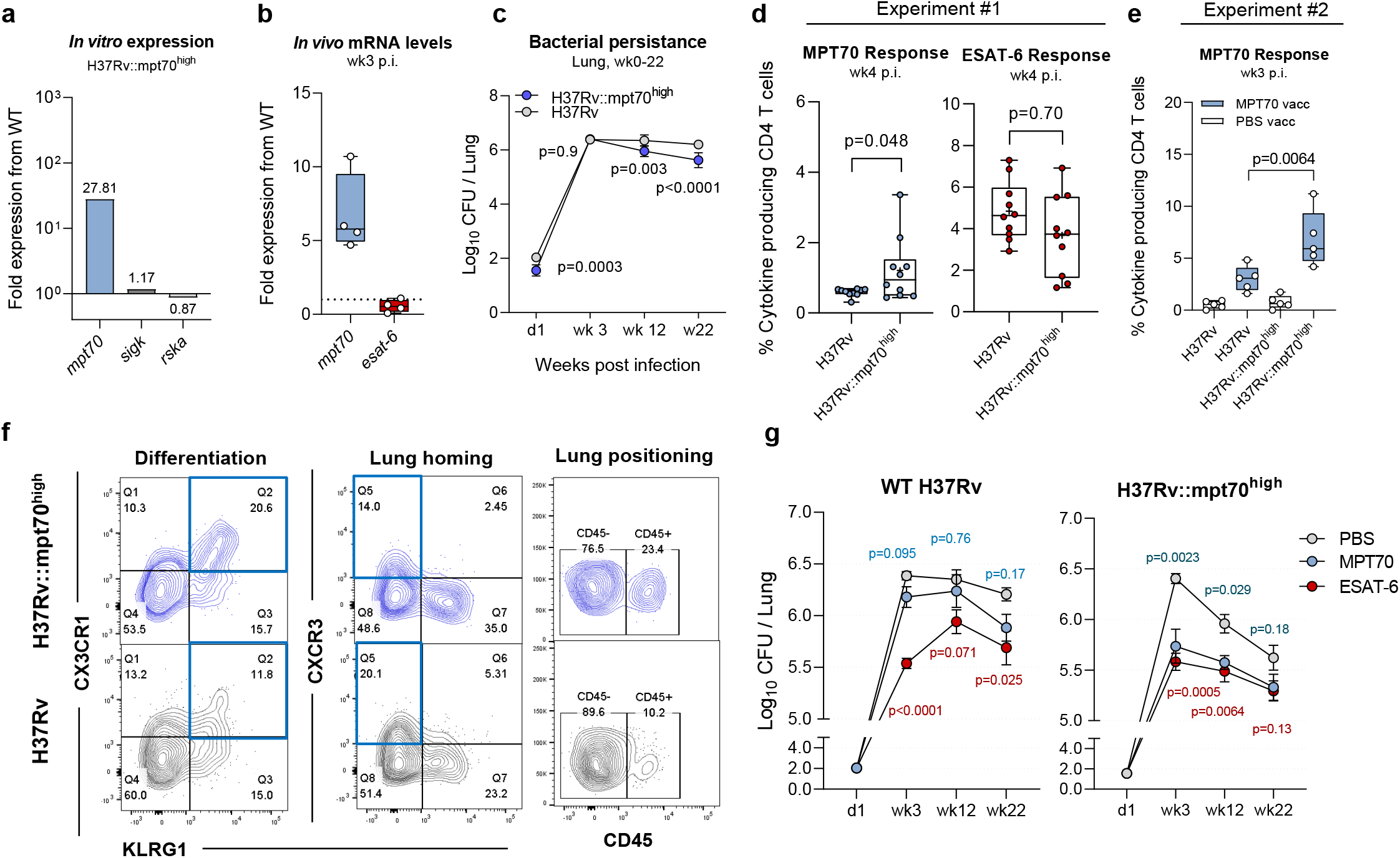
Overexpression of MPT70 accelerates T cell differentiation and improves vaccine efficacy. **(a)** *In vitro* fold gene expression of sigK, rskA, ESAT-6, MPT70, and MPT83 in H37Rv::mpt70^high^ compared to WT H37Rv. All genes were tested in technical duplicates and normalised to esxA expression using primers in **Table 1**. **(b)** *In vivo* fold gene expression of MPT70 and ESAT-6 in lungs ofH37Rv::mpt70^high^ infected mice compared to WT H37Rv infected mice 3 weeks post Mtb challenge (n=5). Genes were analysed in technical duplicates using primers and probes in **Table 2**, normalised to 16s rRNA expression, and shown as fold increase from WT H37Rv. Unpaired t-test, two-tailed. **(c)** Bacterial burden in lungs of PBS-vaccinated mice at day1, week 3, week 12, and week 22 after infection with either H37Rv:: mpt70^high^ or WT H37Rv infection (n=5). Shown as average mean ± SEM. Multiple t-tests with correction for multiple testing using the Holm-Sidak method. **(d)** Frequency of lung MPT70 and ESAT-6-specific CD4 T cells 4 weeks post Mtb infection (n=10). Shown as box plots with whiskers indicating the minimum and maximum values. Mean indicated with ‘+’. Unpaired, two-tailed t-test. **(e)** Frequency of lung MPT70-specific CD4 T cells 3 weeks post Mtb infection in PBS vaccinated (white boxes) and MPT70 vaccinated (blue boxes, n=5). Unpaired, two-tailed t-test. **(f)** Representative concatenated FACS plots (n=10) showing the expression of CX3CR1, CXCR3, KLRG1 or CD45 on MPT70-specific CD4 T cells 4 weeks post H37Rv:: mpt70^high^ infection (blue) or H37Rv infection (grey). Flow Cytometry gating as depicted in Figure S1, using **antibody panel 3. (g)** Bacterial numbers were determined in the lungs of PBS, MPT70 and ESAT-6 vaccinated mice at day 1, week 3, week 12, and week 22 post WT H37Rv infection **(left)** or H37Rv::mpt70^high^ infection **(right)** (n=4-5). One mouse was excluded from the week 12 timepoint (H37Rv::mpt70^high^, MPT70 vaccinated), as the mouse was very sick, had high weight loss, and met the study’s predefined humane endpoints (p-value=0.67, if included). Shown as average mean ± SEM. Statistical differences were assessed using One-Way ANOVA with Tukey’s Multiple Comparison Test.

Taken together, overexpression of MPT70 increased MPT70-vaccine protection substantially, indicating that *in vivo* antigen expression kinetics regulates vaccine protection rather than properties of the antigen as such. This, even though high antigen expression resulted in increased T cell differentiation. Vaccination with highly expressed antigens therefore requires that the vaccine primes T cells of a low differentiation.

## Discussion

CD4 T cell differentiation is a key determinant of protective immunity against Mtb (21, 38, 39) and antigen load is described to influence T cell development (21–23). Importantly, during the course of infection, Mtb adapts to the changing environment of the host and it is poorly described how differential *in vivo* antigen expression influences CD4 T cell responses and vaccine potential. In this study, we compared ESAT-6, which is a well-known constitutively expressed antigen (32, 33), to MPT70 that is expressed at negligible amounts *in vitro* but inducible upon IFN-γ activation (5, 6) or nutrient-deprivation (8) in infected macrophages. After murine aerosol infection, we observed that MPT70 immune recognition was delayed compared to ESAT-6, which correlated with lower initial antigen expression of MPT70. This is in line with a previous genome-wide microarray study after H37Rv infection in Balb/c mice, where MPT70 expression gradually increases from day seven to 28, where it is similar to ESAT-6 (40). Based on this it can be speculated that *in vivo* expression of MPT70 is upregu-lated in response to host adaptive immune responses, which occur around weeks 2-3 in mice (41, 42). This likely places MPT70 in the same functional category as the stress-induced genes encoded by the dormancy survival regulon (DosR) that likewise are upregulated in chronically infected mice and IFN-γ treated macrophages (5, 43).

Given that antigen availability influences T cell quality, we asked how the delayed antigen recognition of MPT70 would impact the CD4 T cell response. In mice infected with WT Mtb Erdman, we observed that MPT70-specific CD4 T cells had a significantly lower differentiation status than ESAT-6 based on the expression of KLRG1, CX3CR1, and T-bet as well as cytokine expression pattern. This is in line with our most recent results, showing that MPT70 is highly immunogenic during late chronic infection, but with an altered T cell phenotype compared to ESAT-6 (29). Having established associations between antigen expression and T cell quality, we investigated whether there was a causal relationship. For this, we utilised a newly described H37Rv strain, which has high *in vitro* expression of MPT70 due to a gene insert of the regulators sigK (Rv0445c) and rskA (Rv0444c) from *M. orygis* (36). Infection with this strain significantly increased the differentiation state of the MPT70-specific CD4 T cells and diminished their ability to enter the infected lung tissue. No differences were observed for ESAT-6 specific CD4 T cells, demonstrating that this was a specific consequence of overexpressing MPT70. These data therefore indicate that the quality of infection-driven T cells is dictated by the *in vivo* antigen expression profile. This is in line with the study by Moguche A *et al*., demonstrating that *in vivo* overexpression of Ag85B significantly increased CD4 T cell differentiation (22). Given that Ag85B expression is downregulated early during infection (32), in contrast to MPT70, these studies collectively suggest CD4 T cell differentiation is a result of the cumulative antigen exposure, and that highly and constitutively expressed antigens would have the highest degree of T cell differentiation.

We finally examined the link between antigen expression and protective capacity in a vaccination setting. Antigens expressed during late-stage Mtb infection have been the focus of multi-stage TB vaccines as they may specifically target bacteria during latency (16, 44–46) and the expression profile of MPT70 could make this an interesting candidate. After vaccination, both MPT70 and ESAT-6 induced robust protection against Mtb Erdman infection, which is in line with other studies reporting high vaccine potential of these two antigens (13, 15, 29, 30). However, despite lower immunogenicity, ESAT-6 vaccination induced the highest protection. Although ESAT-6 and MPT70 are different in both size, epitope pattern, and immunogenicity, this observation prompted us to hypothesise that constitutive *in vivo* antigen expression is optimal for vaccine protection. To investigate this more directly, we compared the effect of immunisation with MPT70 against WT H37Rv or H37Rv::mpt70^high^ where MPT70 is overexpressed. Here we observed that the MPT70-mediated protection was significantly increased against the H37Rv::mpt70^high^ strain, where ESAT-6 and MPT70 performed similarly, compared to the WT H37Rv strain. Together with the T cell analysis, these data suggest that constitutively expressed antigens are superior vaccine antigens, supposedly because of increased antigen “visibility” by the infected macrophages, but due to continuously high antigen presence, the infection-driven T cells are also pushed towards terminal differentiation and decreased functionality. We speculate that this could be a survival strategy by Mtb (47) and show how targeted vaccination with adjuvanted proteins can compensate for this by priming (and maintaining) less differentiated T cells (29).

Importantly, we only investigated the impact of vaccination after single antigen immunisation and it can be speculated that MPT70, and similar antigens, might perform differently in larger fusions proteins. Here, the accelerated adaptive immune responses offered by other antigens, like ESAT-6 (41), may trigger earlier expression of MPT70, which in turn would increase the MPT70 mediated protection against Mtb. Additionally, MPT70 is naturally overexpressed in the animal-adapted *Mycobacterium* strains; *M. orygis*, *M. caprae*, and *M. bovis* (6), which could imply that an MPT70-containing vaccine would be particularly efficacious against pathogens causing TB in livestock.

The course of chronic infection mimicked in the mouse model is in many ways different from the human Mtb infection that can last for years and display distinct features in granuloma structure and environment (48). The findings of this study therefore need to be investigated and validated in human studies, where ESAT-6 and MPT70 could be used as model antigens. There may also be differences in antigen expression levels between clinical Mtb isolates, that are known to display some level of genetic diversity and virulence variability (49) and future studies should extrapolate our results to other relevant isolates. Of note, MPT70 has been described as part of the “core transcriptome” in macrophage phagosomes with conserved expression and regulation across all MTBC isolates (50), suggesting that *in vivo* MPT70 expression will not vary between clinical Mtb isolates. Overexpression of MPT70 however, did seem to impact the bacteria’s overall capability to persist after week three, implicating that abundant MPT70 is not advantageous for Mtb during chronic infection. This has also been seen in a similar study overexpressing Mtb heat-shock proteins (51).

Overall, our study provides new insights into host-pathogen interactions and describes how *in vivo* antigen expression kinetics can regulate T cell functionality and vaccine protection. Data show that high antigen expression drives T cells toward terminal differentiation and that targeted preventive vaccination can counteract this effect. We also demonstrate that highly expressed antigens are optimal vaccine targets and accentuate that T cell differentiation can be used as a new way to identify the most promising antigens. This has implications for rational vaccine design and future pre-clinical and clinical studies could use antigen-specific T cell differentiation as a readily measurable proxy for high *in vivo* antigen expression.

## Author Contributions

HSC, RM, MAB, CAA, and PA conceived and designed the studies. HSC, JYD, FM performed murine TB experiments. HSC and RM analysed and interpreted the data. GJ took part in the supervision and provided intellectual content to the study. IR designed and produced the recombinant proteins including quality control and testing. HSC and RM drafted the manuscript. HSC, MAB, PA, and RM finalised the manuscript. All authors reviewed and commented on the final manuscript.

## Data Sharing Statement

The data that support the findings of this study are available on request from the corresponding author.

## Acknowledgment

We acknowledge the NIH Tetramer Core Facility for provision of I-Ab:ESAT-6_4-17_ and I-A^b^:MPT70_38-52_ and corresponding negative control tetramers I-Ab:hCLIP and the Containment Level 3 laboratory of the Research Institute of the McGill University Health Centre. Thanks to Andréanne Lupien and Sarah Danchuk for technical assistance and fruitful discussion on RNA extractions and qPCR assays.

We thank Vivi Andersen, Ming Liu Olsen, and Camilla Haumann Rasmussen at SSI for their excellent technical assistance. We also gratefully acknowledge the mouse work done by the competent veterinarians and dedicated animal caretakers at Statens Serum Institut.

## Methods and Data Availability

### Mice

Six to eight week old female CB6F1 mice (BALB/c x C57BL/6, Envigo) or C57Bl/6 mice (Jackson Laboratory) were purchased. Mice were randomly assigned to cages of five to eight on the day of arrival. Before initiating the experiment, mice had at least one week of acclimation. Mice were housed in Biosafety Level (BSL) II in individually ventilated cages (Scanbur, Denmark) and had access to nesting material as well as enrichment. During the course of the experiment, mice were fed with irradiated Teklad Global 16% Protein Rodent Diet (Envigo, 2916C) and had access to water ad libitum. On the day of challenge, cages with mice were transferred to BSL-III where they were housed until termination of the experiment.

### Ethics

All experimental protocols were initially reviewed and approved by a local ethical committee at Statens Serum Institut (SSI) and by the Facility Animal Care Committee at the Research Institute of McGill University Health Center (RI-MUHC) (project ID 2015-7656). Experimental procedures were conducted in accordance with the regulations set forward by the Danish Ministry of Justice, Canadian Council of Animal Care (CCAC), and Animal Protection Committees under license permit no. 2019-15-0201-00309 and in compliance with the European Union Directive 2010/63/EU.

### Recombinant Proteins & Immunisations

The following recombinant antigens were expressed and purified as previously described (29): ESAT-6 (Rv3875) or MPT70 (Rv2875). Mice were immunised three times subcutaneously (s.c.) at the base of the tail or neck at two weeks intervals with either recombinant ESAT-6 or MPT70. Recombinant proteins were diluted in Tris-HCL buffer + 9% Trehalose (pH 7.2) and adjuvanted with cationic adjuvant formulation 1 (CAF01^®^) consisting of dimethyldioctadecylammonium (DDA) and trehalose dibehenate (TDB) in a ratio 250 μg DDA per/50 μg TDB (35). As a control, mice were vaccinated with Phosphate Buffered Saline (PBS).

### Mtb Infections and CFU Enumeration

Mtb Erdman (ATCC 35801 / TMC107) was cultured in Difco ^™^ Middlebrook 7H9 (BD) supplemented with 10% BBL ^™^ Middlebrook ADC Enrichment (BD) for two-three weeks using an orbital shaker (~110 rpm, 37°C). Bacteria were harvested in log phase and stored at − 80°C until use. Before used in the experiment the concentration of the bacterial stock was determined by plating in triplicate. For aerosol infections, the vial of Mtb was thawed, sonicated for five minutes, thoroughly suspended with a 27G needle to remove clumps, and mixed in PBS to the desired concentration. Mice were challenged with 0.5 × 10^6^ CFU/mL (around 50-100 CFUs) Mtb Erdman by the aerosol route using a Biaera exposure system controlled via AeroMP software.

The following strains; Mtb Pasteur H37Rv, Pasteur H37Rv::mpt70^high^ (*rskA* and *sigK* of *M. orygis*) (36), and Pasteur H37Rv::Rv (*rskA* and *sigK*of *M. tuberculosis*) (36) were grown in Middlebrook 7H9 medium (Difco Laboratories) supplemented with 0.05% Tween 80 (Sigma-Aldrich), 0.2% glycerol, 10% ADC Enrichment (BD) in a rolling incubator at 37°C. The bacterial cultures were passaged twice and adjusted to an OD_600_ of 0.5, pelleted and resuspended in glycerol, and subsequently frozen at − 80°C. On the day of the experiment, vials were thawed, thoroughly resuspended with a 27G needle, and adjusted to an OD of 0.05 in PBS (approximately 50 CFUs). Mice were challenged with an aerosol infection using a CH Technologies Nose-Only Inhalation Exposure System system with 15 minutes exposure, up to 18 mice per run. Mice euthanised for each experimental time point were in the same aerosol run.

To enumerate bacteria in the lungs of mice after infection, left lobes from individual mice were homogenised with GentleMACS M-tubes (Miltenyi Biotec) in 3 mL MilliQ water containing PANTA^™^ Antibiotic Mixture (BD, cat.no. #245114) or with an Omni Tissue Homogeniser and Hard Tissue Omni Tip^™^ Plastic Homogenising Probes (Omni International) in a 50 mL tube containing 1 mL 7H9 supplemented medium. The homogenate was serially diluted, plated, and grown on 7H11 plates (BD) or 7H10 plates containing PANTA^™^ for approximately 2-3 weeks at 37°C and 5% CO_2_. CFUs were counted, log-transformed to normalise data, and shown as log_10_ CFU per the whole lung. Whenever possible a cutoff of 10 colonies was set to minimise variability and errors due to plating.

### *In vitro* Growth Assay

The growth of Mtb H37Rv, Mtb H37Rv::mpt70^high^ and H37Rv::Rv in 7H9 medium (supplemented as described above) was monitored with a spectrophotometer at OD_600_ in triplicates every 24h for 4 days. The culture flasks were incubated at 37°C under rotating conditions.

### *In vitro and in vivo* Mtb RNA extractions

For *in vitro* RNA extractions, Mtb H37Rv and H37Rv::mpt70^high^ were passaged twice and adjusted to an OD_600_ of 0.2-0.5. The bacteria were pelleted and resuspended in 1 mL TRIzol^™^ Reagent (Invitrogen, Cat. No. 15596026). For *in vivo* RNA extractions the post-caval lung lobes of Mtb infected mice were harvested aseptically and immediately stored at − 80°C in 1 mL RNA later (Qiagen, Cat No./ID: 76106) until further processing. The lung tissues were mechanically disrupted in 1 mL TRIzol^™^ Reagent using Lysing Matrix D or E tubes (MP, SKU: 116913050-CF, SKU: 116914050-CF) and a FastPrep-24^™^ bead beater (MP, SKU: 116004500). The grinded lung tissues were stored at − 80°C until further processing. On the day of RNA extraction, the lung tissues were bead beated with 0.1 mm Zirconia/Silica beads three times at 6.5 m/s for 30 seconds with 3 minutes rest on ice in between runs. The beads were pelleted by centrifuging at 12,000 g for 1 minute and the TRIzol layer moved to a fresh tube containing chloroform isoamyl alcohol (24:1). After centrifuging at 12,000 g for 15 minutes at 4°C, the top aqueous phase was transferred and precipitated with 3M sodium acetate and isopropanol for at least 2 hours or overnight at − 20°C. The pellet was washed twice in ethanol, air-dried, and resuspended in RNAse free water. A cleanup step was performed with RNA Easy Mini Kit (Qiagen), followed by a minimum of three DNAse treatments (Ambion, Cat. No. AM1907). The RNA purity and concentration were measured by spectrophotometry (Tecan Infinite M200 Pro plate reader) or using the RNA Qubit Assay (Invitrogen^™^, Cat.no. Q32852). RNA samples were checked for remaining genomic DNA by PCR using the SigA primers. A total of 300-1000ng RNA was reverse transcribed (Thermo Scientific, Cat.no. K1621); a minus reverse transcriptase control was included for every sample.

### Real-time qPCR and Gene Expression Analysis

To determine *in vitro* mRNA levels of MPT70, SigK, RskA, and ESAT-6, we performed an RT-qPCR using Maxima SYBR green kit (Thermo Scientific, Cat.no.K0223) with the use of the following primers **(Table 1)**. mRNA levels were normalised to EsxA and fold gene expression from Mtb H37Rv was plotted as 2^-^ (ΔCt)

**Table 1.**
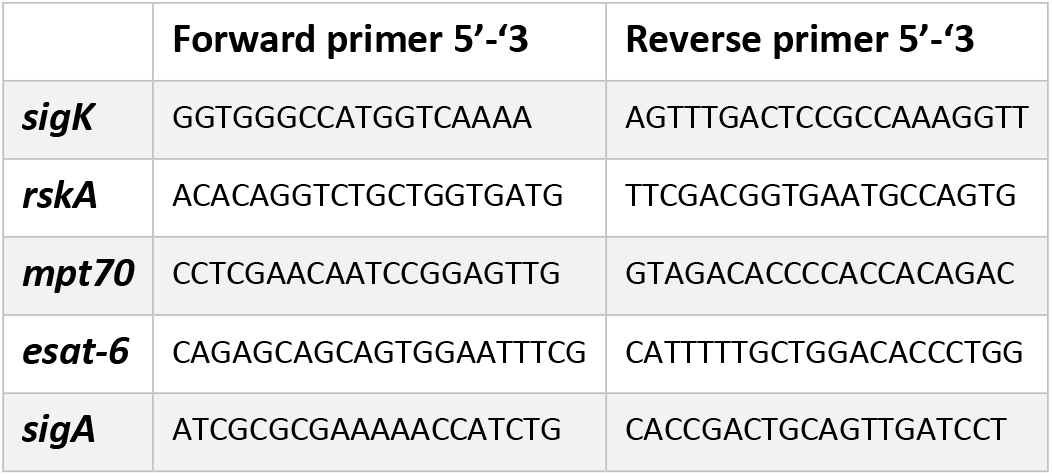
Primers used for *in vitro* gene expression analyses.

*In vivo* mRNA levels were measured with qRT-PCR using dual-labelled probes (Eurofins) **(Table 2)**. All probes and primers were diluted to a final concentration of 250 nM (probes) and 900 nM (primers) respectively, and mixed with either iTaq Universal Probe Supermix (Biorad, cat. no. 1725130) or SsoAdvanced Universal Probes Supermix (Biorad, Cat. no. 1725281). All cDNA samples were prediluted 10x in DEPC-treated water and used in a final dilution of 1:40 in the reaction. Thermal cycling protocol was programmed according to the manufacturer’s instructions for low abundant targets (95° 30 seconds; 95° 10 seconds; 60° 1 minute, 45 cycles). For gene expression analysis throughout Mtb Erdman infection **(Figure 1a)**, average Cq values for each sample were normalised to 16s rRNA and shown as relative mRNA levels (2^-(ΔCq)^). The fold gene expression of *mpt70* and *esat-6* in H37Rv::mpt70^high^ from WT H37Rv were calculated with the 2^-*ΔΔCT*^ method with normalisation to 16s rRNA **(Figure 4b)**. For every run, a no template control, negative control (naïve mouse), and positive controls (genomic DNA of H37Rv and BCG) were included.

**Table 2.**
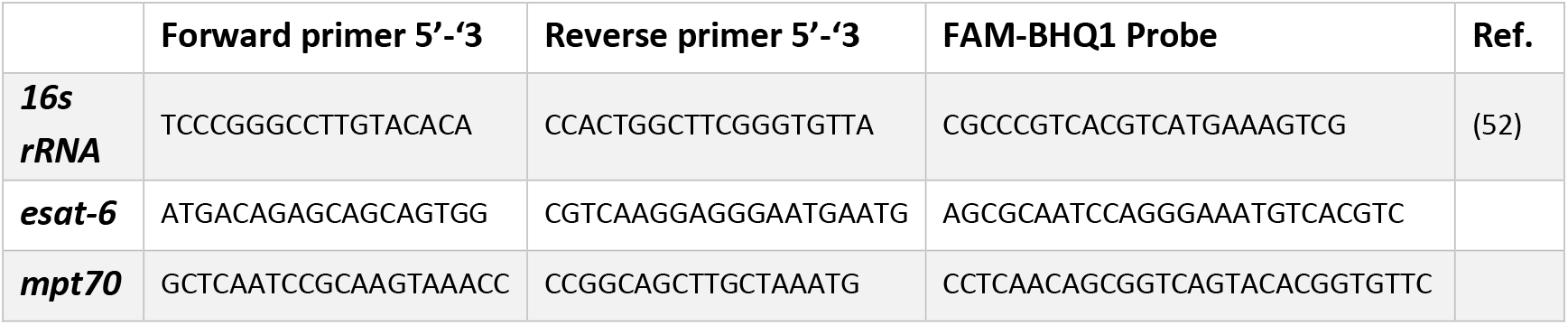
Primers and probes used for *in vivo* gene expression analyses.

### *In vivo* Labelling of Intravascular CD4 T cells

Mice were anesthetised with isoflurane and injected intravenously with 2.5-5.0 μg fluorescein isothiocyanate (FITC) labelled CD45 antibody (BD Pharmingen, clone 104; 553772) diluted in 100-250 μl PBS. Three minutes after injection, mice were euthanised by cervical dislocation, and organs aseptically harvested for further processing as described below.

### Preparation of Single-Cell Suspensions

Spleens or lungs were aseptically harvested from euthanised mice. Lungs were first homogenised in Gentle MACS tubes C (Miltenyi Biotec) or chopped into small pieces using scalpels, followed by 1 hour of collagenase-digestion (Sigma Aldrich; C5138) at 37°C, 5% CO_2_. The lung homogenate and spleens were forced through 70-100 μm cell strainers (BD Biosciences) with the stopper from a 5 mL syringe (BD) and washed twice with cold RPMI medium (Gibco; RPMI-1640) by centrifuging 5 minutes at 1800 rpm. A red blood cell lysis step was performed in between washes (Roche, cat. no. 11814389001). Cells were finally resuspended in enriched RPMI medium (RPMI-1640, 10% heat-inactivated FCS (Biochrom Gmbh), 10 mM Hepes (Invitrogen), 2 mM L-Glutamine (Invitrogen), 1 mM Natriumpyruvate (Invitrogen), 1× Non-essential amino acids (MP Biomedicals, LLC), 5×10^5^ M 2-mercaptoethanol (Sigma-Aldrich) and Penicillin-Streptomycin (Gibco)). Cells were counted using an automatic Nucleocounter^TM^ (Chemotec) and adjusted to 2×10^5^ cells/well for ELISA and 1-2×10^6^ cells/well for flow cytometry.

### Design of an MHC-II Tetramer specific for MPT70

Splenocytes of MPT70-vaccinated mice were restimulated in the presence of overlapping 15-mer peptides and two murine epitopes were identified (29). The recognised peptides corresponded to amino acid (aa) location 37-53 and 93-109 in MPT70. These epitopes were further epitope mapped in vaccinated mice with peptides varying of one aa in length to identify the minimal core epitope which is the minimal number of aa necessary for T cell recognition (**Figure S2a**). We found EYAAANPTGPA and FAPTNAAF as the core epitopes, but as FAPTNAAF was not highly recognised by Mtb infected mice (data not shown), it was not further characterised. As the optimal peptide length for an MHC-II tetramer may vary between 11-16 aa, different peptide lengths extending the core epitope were tested with no big difference in response magnitude as long as the sequence “YAAANPTGP” were present (**Figure S2b**). Based on these results we designed a tetramer specific for I-A^b^:MPT70_38-52_ (CAEYAAANPTGPAS). This epitope has previously been identified in MPB70 DNA-immunised C57Bl/6 H-2^b^ and B6D2 (F1) H-2d^b^ **mice** in an ELISPOT assay with no humoral response detected in mice of haplotype H-2^d^ (53).

### MHC-II Tetramer Staining

Tetramers (MPT70 ^38-52^:I-Ab, ESAT-6^4-17^:I-Ab) conjugated to BV421 or PE and corresponding negative controls (hCLIP:I-Ab) were provided by the NIH tetramer core facility (Atlanta, USA). MHC-II tetramers were titrated and tested for optimal staining conditions before the experiment. Single-cell suspensions were stained with tetramers diluted 1:50 in FACS buffer (PBS+1%FCS) containing 1:200 Fc-block (anti-CD16/CD32) for 30 minutes at 37°C. The MPT70^38-52^ MHC-II tetramer was specifically developed for this study **(Figure S2)**.

### *In Vitro* Re-stimulation and Intracellular Cytokine Staining

For intracellularly cytokine staining (ICS), cells were restimulated with 2 μg/mL antigen or medium in the presence of 1 μg/ml anti-CD28 (clone 37.51) and anti-CD49d (clone 9C10-MFR4.B) in 96V-bottom TCT microtiter plates (Corning; 3894) for 1 hour at 37°C and 5% CO_2_. Restimulation with ionomycin in conjunction with phorbol myristate acetate (PMA) was included as a positive control. Subsequently, 10 μg/mL Brefeldin A was added to each well (Sigma Aldrich; B7651-5mg) and followed by another 56 hours incubation at 37°C, 5% CO_2_, after which cells were kept at 4°C until staining or immediately surface stained and fixed.

Prior to staining, cells were washed with FACS buffer and subsequently stained surface markers diluted in brilliant stain buffer (BD Horizon; 566349) using antibodies indicated in **Panel 1, 2, and 3**. Fixation and permeabilisation were performed using the Fixation/Permeabilization Solution Kit (BD Cy-tofix/Cytoperm; 554714) or Foxp3 / Transcription Factor Staining Buffer Set (eBioscience^™^; 00-5523-00) as per manufacturer’s instructions followed by intracellular staining (ICS) with anti-IFN-γ, anti-IL-2, anti-TNF-α, and anti-IL17A and/or transcription factor staining with anti-T-bet. Fluorescence minus one controls were performed for CD3, CD44, KLRG1, PD-1, CXCR3, CX3CR1, IL-2, IL-17, IFN-γ, and T-bet on pooled cells to set boundaries gates for surface-, intracellular-and transcription factor markers. Gating strategies for defining tetramer-positive CD4 T cells and cytokine-producing CD4 T cells are exemplified in **Figure S3** (Tetramer) and **Figure S1** (ICS).

**Antibody Panel 1:**

**Antibodies used for tetramer characterisation in** Figure 2.

| ANTIBODY | FLOUREPHORE | SUPPLIER | CLONE | DILUTION | CAT NO |
| --- | --- | --- | --- | --- | --- |
| CD3 | Bv650 | Biolegend | 17A2 | 1:100 | 100229 |
| CD4 | Bv510 | Biolegend | RM4-5 | 1:400 | 100559 |
| VIABILITY | eflour780 | eBioscience |  | 1:1000 |  |
| CXCR3 | PerCpCy5.5 | eBioscience | CXCR3-173 | 1:100 | 45-1831-82 |
| CX3CR1 | Bv785 | Biolegend | SAO11F11 | 1:100 | 149029 |
| CD44 | Alx700 | Biolegend | IM7 | 1:200 | 103026 |
| KLRG1 | Bv711 | Biolegend | 2F1 | 1:100 | 138427 |
| CD45 (*in vivo*) | FITC | BD Bioscience | 104 | 1:50 | 553772 |
| T-BET | PE-Cy7 | eBioscience | 4B10 | 1:100 | 25-5825-82 |
| HCLIP CON | Bv421 | NIH | - | 1:50 | - |
| MPT70 TET | Bv421 | NIH | - | 1:50 | - |
| HCLIP CON | PE | NIH | - | 1:50 | - |
| E6 TET | PE | NIH | - | 1:50 | - |

**Antibody Panel 2:**

Antibodies used for T cell characterisation in **Figures 1, 2, and 3**.

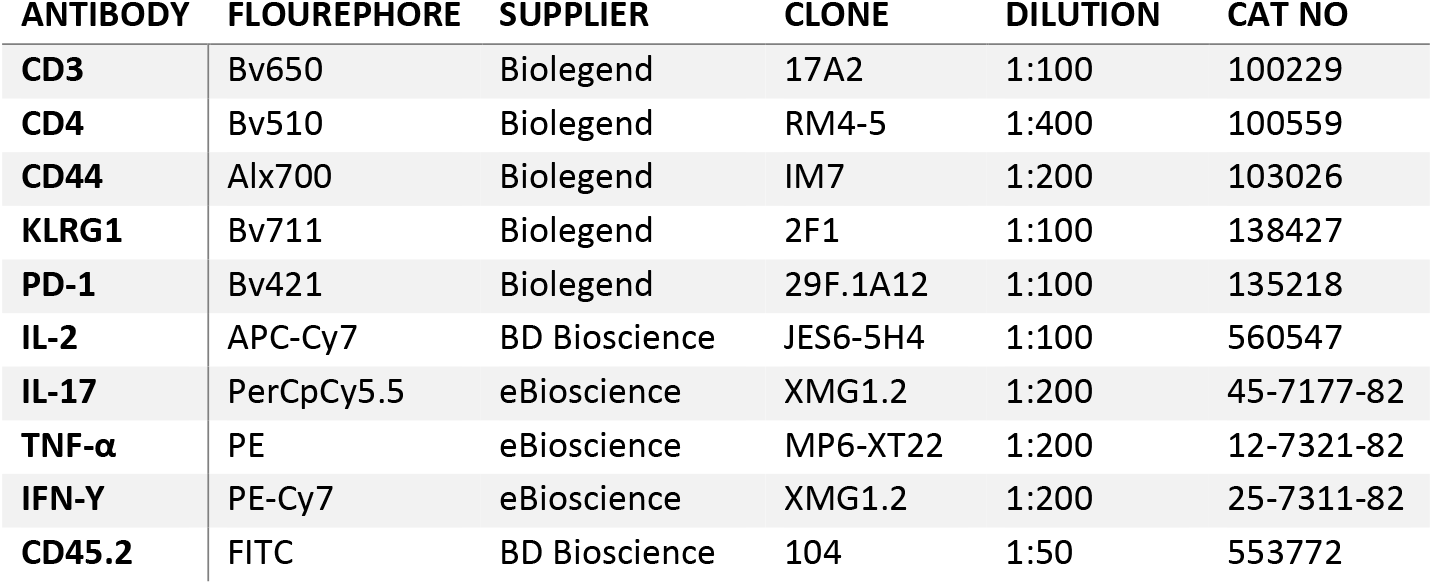

**Antibody Panel 2:** Antibodies used for T cell characterisation in **Figures 1, 2, and 3.**
| ANTIBODY | FLOUREPHORE | SUPPLIER | CLONE | DILUTION | CAT NO |
| --- | --- | --- | --- | --- | --- |
| CD3 | Bv650 | Biolegend | 17A2 | 1:100 | 100229 |
| CD4 | Bv510 | Biolegend | RM4-5 | 1:400 | 100559 |
| CD44 | Alx700 | Biolegend | IM7 | 1:200 | 103026 |
| KLRG1 | Bv711 | Biolegend | 2F1 | 1:100 | 138427 |
| PD-1 | Bv421 | Biolegend | 29F.1A12 | 1:100 | 135218 |
| IL-2 | APC-Cy7 | BD Bioscience | JES6-5H4 | 1:100 | 560547 |
| IL-17 | PerCpCy5.5 | eBioscience | XMG1.2 | 1:200 | 45-7177-82 |
| TNF-α | PE | eBioscience | MP6-XT22 | 1:200 | 12-7321-82 |
| IFN-Y | PE-Cy7 | eBioscience | XMG1.2 | 1:200 | 25-7311-82 |
| CD45.2 | FITC | BD Bioscience | 104 | 1:50 | 553772 |

**Antibody Panel 3:**

**Antibodies used for T cell characterisation in** Figure 4.

**Antibody Panel 3:** Antibodies used for T cell characterisation in **Figure 4.**
| ANTIBODY | FLOUREPHORE | SUPPLIER | CLONE | DILUTION | CAT NO. |
| --- | --- | --- | --- | --- | --- |
| VIABILITY | 440UV | BD Horizon™ | - | 1:1000 | 566332 |
| CD8A | APC-H7 | BD Pharmingen™ | 53-6.7 | 1:100 | 560182 |
| CD4 | PE | BD Pharmingen™ | GK1.5 | 1:400 | 557308 |
| CD45 | FITC | BD Pharmingen™ | 30-F11 | 1:10 | 553080 |
| CD44 | BUV395 | BD OptiBuild™ | IM7 | 1:100 | 740215 |
| KLRG1 | BB700 | BD OptiBuild™ | 2F1 | 1:100 | 742199 |
| PD-1 (CD279) | APC-R700 | BD Horizon™ | J43 | 1:100 | 565815 |
| CXCR3 (CD183) | BUV737 | BD OptiBuild™ | CXCR3-173 | 1:100 | 741895 |
| CX3CR1 | PE/Dazzle™ 594 | Biolegend | SA011F11 | 1:100 | 149014 |
| CD3E | V500 | BD Horizon™ | 500A2 | 1:50 | 560771 |
| TNF-α | PE-Cy™7 | BD Pharmingen™ | MP6-XT22 | 1:200 | 557644 |
| IFN-Y | Bv421 | BD Horizon™ | XMG1.2 | 1:100 | 563376 |
| IL-17A | BV786 | BD Horizon™ | TC11-18H10 | 1:100 | 564171 |
| IL-2 | APC | eBioscience | JES6-5H4 | 1:100 | 17-7021-82 |

### IFN-y Sandwich ELISA

Splenocytes or lung cells were adjusted to a cell concentration of 2×10^5^ cells/well and restimulated in the presence of recombinant protein or peptides in round-bottom plates for 3 days as previously described (29). A sandwich ELISA was performed on the culture supernatants to determine the concentration of total IFN-γ. In brief, microtiter plates were coated with primary IFN-γ antibody, blocked with 2% skimmed milk, and incubated overnight with pre-diluted supernatants. IFN-γ was detected with a secondary IFN-γ antibody, followed by an HRP-conjugated antibody and the reaction was developed using TMB substrate (TMB Plus; Kementec). Plates were read at 450 nm with 620 nm background correction using an ELISA reader (Tecan Sunrise).

### Statistical Analyses

Cells from the murine studies were analysed using a BD LRSFortessa or BD LRSFortessa X20 and the FSC files were afterwards manually gated with FlowJo v10 (Tree Star). The graphical visualisations were done using GraphPad Prism v8.3.0. The type of statistical test performed together with the exact p-value is indicated in the individual figure legends. A p-value below 0.05 was considered significant.

## Supplementary Figure Legends

**Figure S1: Gating strategy for antigen-specific CD4 T cells after intracellular staining (ICS).** Spleens and lungs were harvested from mice and prepared as single-cell suspensions. Shown as representative gating from sample WT H37Rv, ESAT-6-specific T cells 3 weeks post Mtb infection using **antibody panel 3**. Cells were gated as singlets, lymphocytes, and CD3^+^CD4^+^ or CD8^+^CD4 positive T cells. CD44^high^ CD4 T cells were analysed for their intracellular production of IFN-γ, TNF-α, IL-2, and IL-17A. The frequency of antigen-specific CD4 T cells was determined by a make or gate for IFN-γ, TNF-α, IL-2, and IL-17A (i.e. T cells can produce one or more of the cytokines). T cell differentiation degrees were analysed with a combination gate for IFN-γ, TNF-α, and IL-2, characterising the cytokine subsets of T cells. The frequency of CD45^+^, KLRG1, PD-1, CXCR3, and CX3CR1 was assessed on antigen-specific T cells. Fluorescence minus one (FMO) controls were used to set boundaries gates for CD44, KLRG1, PD-1, CXCR3, and CX3CR1.

**Figure S2: Epitope mapping and design of an MPT70 tetramer. (a)** Splenocytes of MPT70-vaccinated mice were *in vitro* restimulated with overlapping peptides of 15 amino acids in length for 3 days (n=4). The amount of IFN-γ was measured in the culture supernatant. The dominant epitope required for binding is highlighted in bold blue text and the predicted core epitope in bold black text. **(b, left)** The minimal epitope of the 38-53 sequence of MPT70 was investigated with varying lengths of peptides in MPT70 vaccinated mice, 20 weeks post-infection (n=4). **(b, right)** Comparison of the response to medium, the chosen 38-53 epitope, and recombinant MPT70. Same data as in left b panel.

**Figure S3: Phenotyping of MPT70_38-52_ and ESAT-6_4-17_ CD4 T cells during Mtb infection.** Gating strategy for tetramer-positive CD4 T cells. Lung cells of vaccinated and infected mice were prepared as single-cell suspensions and analysed by flow cytometry. Shown as representative gating for tetramer-positive CD4 T cells exemplified with saline mouse A6 infected for 16 weeks using **antibody panel 1**. Cells were gated as singlets and lymphocytes. Viable CD3^+^ CD4^+^ CD44^high^ T cells were stained with either I-A^b^:MPT70_38-52_ and I-Ab:ESAT-6_4-47_ tetramer. A corresponding control tetramer, hClip, was included. Tetramer positive CD4 T cells were further characterised for their expression of KLRG1, T-bet, and CXCR3. Fluorescence minus one (FMO) controls were used to set boundaries gates for KLRG1, T-bet, and CXCR3.

**Figure S4: Long-term vaccine impact of ESAT-6 and MPT70 during Mtb infection.** CB6F1 mice were vaccinated with MPT70, ESAT-6, or saline three times and challenged with Mtb Erdman 6 weeks post 3^rd^ immunisation. **(a)** Percentage of KLRG1^+^PD-1^-^ of MPT70 or ESAT-6-specific CD4 T cells in vaccinated and saline mice 3 and 20 weeks post Mtb infection (n=4). Shown as box plots with whiskers indicating the minimum and maximum values. **(b)** The bacterial burdens were determined in the lungs of saline and vaccinated mice 19 or 20 weeks post Mtb infection (n=28). The graph represents four individual experiments. One-Way ANOVA with Tukey’s multiple comparison test.

**Figure S5: Characterisation of the modified H37Rv::mpt70^hlgh^ strain**. **(a)** Relative mRNA levels of MPT70 and ESAT-6 in lungs of WT H37Rv and H37Rv::mpt70^high^ infected mice 3 weeks post aerosol Mtb challenge (n=5). mRNA levels were normalised to 16s rRNA. Shown as box plots with whiskers indicating the minimum and maximum values. Paired t-test, two-tailed. **(b)** *In vitro* growth of WT H37RV, H37Rv::mpt70^high^ (rskA and sigK insert of *M.orygis* origin), and H37Rv::Rv (rskA and sigK insert of Mtb origin). Strains were grown in 7H9 medium for 4 days and the OD_600_ was measured every 24 hours (n=3). Shown as average mean ± SD. Multiple t-tests with correction for multiple tests using the Holm-Sidak method. **(c)** KLRG1^+^CX3CR1^+^ expressing MPT70 specific CD4 T cells in PBS vaccinated and MPT70-vaccinated mice 3 weeks post WT H37RV and H37Rv::mpt70^high^ infection (n=5). Shown as individual mice and the average mean. **(d)** KLRG1^+^CX3CR1^+^ expressing MPT70 and ESAT-6 specific CD4 T cells in PBS vaccinated mice, 3-4 weeks post WT H37RV and H37Rv::mpt70^high^ infection (n=5-10). Shown as box plots with whiskers indicating the minimum and maximum values. Two independent experiments. Unpaired, two-tailed t-test.

## References

1. Anonymous. 2020. Global tuberculosis report 2020. World Health Organization, Geneva.

2. Gautam US, Mehra S, Kaushal D. 2015. In-Vivo Gene Signatures of Mycobacterium tuberculosis in C3HeB/FeJ Mice. PLoS One 10:e0135208.

3. Karakousis PC, Yoshimatsu T, Lamichhane G, Woolwine SC, Nuermberger EL, Grosset J, Bishai WR. 2004. Dormancy phenotype displayed by extracellular Mycobacterium tuberculosis within artificial granulomas in mice. J Exp Med 200:647–57.

4. Sharma D, Bose A, Shakila H, Das TK, Tyagi JS, Ramanathan VD. 2006. Expression of mycobacterial cell division protein, FtsZ, and dormancy proteins, DevR and Acr, within lung granulomas throughout guinea pig infection. FEMS Immunol Med Microbiol 48:329–36.

5. Schnappinger D, Ehrt S, Voskuil MI, Liu Y, Mangan JA, Monahan IM, Dolganov G, Efron B, Butcher PD, Nathan C, Schoolnik GK. 2003. Transcriptional Adaptation of Mycobacterium tuberculosis within Macrophages: Insights into the Phagosomal Environment. J Exp Med 198:693–704.

6. Said-Salim B, Mostowy S, Kristof AS, Behr MA. 2006. Mutations in Mycobacterium tuberculosis Rv0444c, the gene encoding anti-SigK, explain high level expression of MPB70 and MPB83 in Mycobacterium bovis. Mol Microbiol 62:1251–63.

7. Harboe M, Nagai S. 1984. MPB70, a unique antigen of Mycobacterium bovis BCG. Am Rev Respir Dis 129:444–52.

8. Betts JC, Lukey PT, Robb LC, McAdam RA, Duncan K. 2002. Evaluation of a nutrient starvation model of Mycobacterium tuberculosis persistence by gene and protein expression profiling. Mol Microbiol 43:717–31.

9. Fifis T, Corner LA, Rothel JS, Wood PR. 1994. Cellular and humoral immune responses of cattle to purified Mycobacterium bovis antigens. Scand J Immunol 39:267–74.

10. Haslov K, Andersen AB, Bentzon MW. 1987. Biological activity in sensitized guinea pigs of MPB 70, a protein specific for some strains of Mycobacterium bovis BCG. Scand J Immunol 26:445–54.

11. Wiker HG. 2009. MPB70 and MPB83--major antigens of Mycobacterium bovis. Scand J Immunol 69:492–9.

12. Ireton GC, Greenwald R, Liang H, Esfandiari J, Lyashchenko KP, Reed SG. 2010. Identification of Mycobacterium tuberculosis antigens of high serodiagnostic value. Clin Vaccine Immunol 17:1539–47.

13. Bertholet S, Ireton GC, Kahn M, Guderian J, Mohamath R, Stride N, Laughlin EM, Baldwin SL, Vedvick TS, Coler RN, Reed SG. 2008. Identification of human T cell antigens for the development of vaccines against Mycobacterium tuberculosis. J Immunol 181:7948–57.

14. Windish HP, Duthie MS, Misquith A, Ireton G, Lucas E, Laurance JD, Bailor RH, Coler RN, Reed SG. 2011. Protection of mice from Mycobacterium tuberculosis by ID87/GLA-SE, a novel tuberculosis subunit vaccine candidate. Vaccine 29:7842–8.

15. Orr MT, Ireton GC, Beebe EA, Huang PW, Reese VA, Argilla D, Coler RN, Reed SG. 2014. Immune subdominant antigens as vaccine candidates against Mycobacterium tuberculosis. J Immunol 193:2911–8.

16. Ma J, Teng X, Wang X, Fan X, Wu Y, Tian M, Zhou Z, Li L. 2017. A Multistage Subunit Vaccine Effectively Protects Mice Against Primary Progressive Tuberculosis, Latency and Reactivation. EBioMedicine 22:143–154.

17. Caruso AM, Serbina N, Klein E, Triebold K, Bloom BR, Flynn JL. 1999. Mice deficient in CD4 T cells have only transiently diminished levels of IFN-gamma, yet succumb to tuberculosis. J Immunol 162:5407–16.

18. Lawn SD, Myer L, Edwards D, Bekker LG, Wood R. 2009. Short-term and long-term risk of tuberculosis associated with CD4 cell recovery during antiretroviral therapy in South Africa. AIDS 23:1717–25.

19. Lin PL, Rutledge T, Green AM, Bigbee M, Fuhrman C, Klein E, Flynn JL. 2012. CD4 T cell depletion exacerbates acute Mycobacterium tuberculosis while reactivation of latent infection is dependent on severity of tissue depletion in cynomolgus macaques. AIDS Res Hum Retroviruses 28:1693–702.

20. Scanga CA, Mohan VP, Yu K, Joseph H, Tanaka K, Chan J, Flynn JL. 2000. Depletion of CD4(+) T cells causes reactivation of murine persistent tuberculosis despite continued expression of interferon gamma and nitric oxide synthase 2. J Exp Med 192:347–58.

21. Day CL, Abrahams DA, Lerumo L, Janse van Rensburg E, Stone L, O’Rie T, Pienaar B, de Kock M, Kaplan G, Mahomed H, Dheda K, Hanekom WA. 2011. Functional capacity of Mycobacterium tuberculosis-specific T cell responses in humans is associated with mycobacterial load. J Immunol 187:2222–32.

22. Moguche AO, Musvosvi M, Penn-Nicholson A, Plumlee CR, Mearns H, Geldenhuys H, Smit E, Abrahams D, Rozot V, Dintwe O, Hoff ST, Kromann I, Ruhwald M, Bang P, Larson RP, Shafiani S, Ma S, Sherman DR, Sette A, Lindestam Arlehamn CS, McKinney DM, Maecker H, Hanekom WA, Hatherill M, Andersen P, Scriba TJ, Urdahl KB. 2017. Antigen Availability Shapes T Cell Differentiation and Function during Tuberculosis. Cell Host Microbe 21:695–706 e5.

23. Han S, Asoyan A, Rabenstein H, Nakano N, Obst R. 2010. Role of antigen persistence and dose for CD4+ T-cell exhaustion and recovery. Proc Natl Acad Sci U S A 107:20453–8.

24. Sakai S, Kauffman KD, Schenkel JM, McBerry CC, Mayer-Barber KD, Masopust D, Barber DL. 2014. Cutting edge: control of Mycobacterium tuberculosis infection by a subset of lung parenchyma-homing CD4 T cells. J Immunol 192:2965–9.

25. Sallin MA, Sakai S, Kauffman KD, Young HA, Zhu J, Barber DL. 2017. Th1 Differentiation Drives the Accumulation of Intravascular, Non-protective CD4 T Cells during Tuberculosis. Cell Rep 18:3091–3104.

26. Chakravarty SD, Xu J, Lu B, Gerard C, Flynn J, Chan J. 2007. The chemokine receptor CXCR3 attenuates the control of chronic Mycobacterium tuberculosis infection in BALB/c mice. J Immunol 178:1723–35.

27. Hall JD, Kurtz SL, Rigel NW, Gunn BM, Taft-Benz S, Morrison JP, Fong AM, Patel DD, Braunstein M, Kawula TH. 2009. The impact of chemokine receptor CX3CR1 deficiency during respiratory infections with Mycobacterium tuberculosis or Francisella tularensis. Clin Exp Immunol 156:278–84.

28. Seiler P, Aichele P, Bandermann S, Hauser AE, Lu B, Gerard NP, Gerard C, Ehlers S, Mollenkopf HJ, Kaufmann SH. 2003. Early granuloma formation after aerosol Mycobacterium tuberculosis infection is regulated by neutrophils via CXCR3-signaling chemokines. Eur J Immunol 33:2676–86.

29. Clemmensen HS, Knudsen NPH, Billeskov R, Rosenkrands I, Jungersen G, Aagaard C, Andersen P, Mortensen R. 2020. Rescuing ESAT-6 Specific CD4 T Cells From Terminal Differentiation Is Critical for Long-Term Control of Murine Mtb Infection. Front Immunol 11:585359.

30. Hoang T, Aagaard C, Dietrich J, Cassidy JP, Dolganov G, Schoolnik GK, Lundberg CV, Agger EM, Andersen P. 2013. ESAT-6 (EsxA) and TB10.4 (EsxH) based vaccines for pre-and post-exposure tuberculosis vaccination. PLoS One 8:e80579.

31. Reiley WW, Calayag MD, Wittmer ST, Huntington JL, Pearl JE, Fountain JJ, Martino CA, Roberts AD, Cooper AM, Winslow GM, Woodland DL. 2008. ESAT-6-specific CD4 T cell responses to aerosol Mycobacterium tuberculosis infection are initiated in the mediastinal lymph nodes. Proc Natl Acad Sci U S A 105:10961–6.

32. Rogerson BJ, Jung YJ, LaCourse R, Ryan L, Enright N, North RJ. 2006. Expression levels of Mycobacterium tuberculosis antigen-encoding genes versus production levels of antigen-specific T cells during stationary level lung infection in mice. Immunology 118:195–201.

33. Shi L, North R, Gennaro ML. 2004. Effect of growth state on transcription levels of genes encoding major secreted antigens of Mycobacterium tuberculosis in the mouse lung. Infect Immun 72:2420–4.

34. Aagaard C, Knudsen NPH, Sohn I, Izzo AA, Kim H, Kristiansen EH, Lindenstrom T, Agger EM, Rasmussen M, Shin SJ, Rosenkrands I, Andersen P, Mortensen R. 2020. Immunization with Mycobacterium tuberculosis-Specific Antigens Bypasses T Cell Differentiation from Prior Bacillus Calmette-Guerin\ Vaccination and Improves Protection in Mice. J Immunol doi:10.4049/jimmunol.2000563.

35. Agger EM, Rosenkrands I, Hansen J, Brahimi K, Vandahl BS, Aagaard C, Werninghaus K, Kirschning C, Lang R, Christensen D, Theisen M, Follmann F, Andersen P. 2008. Cationic liposomes formulated with synthetic mycobacterial cordfactor (CAF01): a versatile adjuvant for vaccines with different immunological requirements. PLoS One 3:e3116.

36. Veyrier FJ, Nieves C, Lefrancois LH, Trigui H, Vincent AT, Behr MA. 2020. RskA Is a Dual Function Activator-Inhibitor That Controls SigK Activity Across Distinct Bacterial Genera. Front Microbiol 11:558166.

37. Hoft SG, Sallin MA, Kauffman KD, Sakai S, Ganusov VV, Barber DL. 2019. The Rate of CD4 T Cell Entry into the Lungs during Mycobacterium tuberculosis Infection Is Determined by Partial and Opposing Effects of Multiple Chemokine Receptors. Infect Immun 87.

38. Lewinsohn DA, Lewinsohn DM, Scriba TJ. 2017. Polyfunctional CD4+ T Cells As Targets for Tuberculosis Vaccination. 8.

39. Petruccioli E, Petrone L, Vanini V, Sampaolesi A, Gualano G, Girardi E, Palmieri F, Goletti D. 2013. IFNgamma/TNFalpha specific-cells and effector memory phenotype associate with active tuberculosis. J Infect 66:475–86.

40. Talaat AM, Lyons R, Howard ST, Johnston SA. 2004. The temporal expression profile of <em>Mycobacterium tuberculosis</em> infection in mice. 101:4602–4607.

41. Woodworth JS, Cohen SB, Moguche AO, Plumlee CR, Agger EM, Urdahl KB, Andersen P. 2017. Subunit vaccine H56/CAF01 induces a population of circulating CD4 T cells that traffic into the Mycobacterium tuberculosis-infected lung. Mucosal Immunol 10:555–564.

42. Chackerian AA, Alt JM, Perera TV, Dascher CC, Behar SM. 2002. Dissemination of Mycobacterium tuberculosis is influenced by host factors and precedes the initiation of T-cell immunity. Infect Immun 70:4501–9.

43. Shi L, Jung YJ, Tyagi S, Gennaro ML, North RJ. 2003. Expression of Th1-mediated immunity in mouse lungs induces a Mycobacterium tuberculosis transcription pattern characteristic of nonreplicating persistence. Proc Natl Acad Sci U S A 100:241–6.

44. Dey B, Jain R, Gupta UD, Katoch VM, Ramanathan VD, Tyagi AK. 2011. A booster vaccine expressing a latency-associated antigen augments BCG induced immunity and confers enhanced protection against tuberculosis. PLoS One 6:e23360.

45. Khademi F, Derakhshan M, Yousefi-Avarvand A, Tafaghodi M, Soleimanpour S. 2018. Multi-stage subunit vaccines against Mycobacterium tuberculosis: an alternative to the BCG vaccine or a BCG-prime boost? Expert Rev Vaccines 17:31–44.

46. Aagaard C, Hoang T, Dietrich J, Cardona PJ, Izzo A, Dolganov G, Schoolnik GK, Cassidy JP, Billeskov R, Andersen P. 2011. A multistage tuberculosis vaccine that confers efficient protection before and after exposure. Nat Med 17:189–94.

47. Comas I, Chakravartti J, Small PM, Galagan J, Niemann S, Kremer K, Ernst JD, Gagneux S. 2010. Human T cell epitopes of Mycobacterium tuberculosis are evolutionarily hyperconserved. Nat Genet 42:498–503.

48. Kramnik I, Beamer G. 2016. Mouse models of human TB pathology: roles in the analysis of necrosis and the development of host-directed therapies. Semin Immunopathol 38:221–37.

49. Peters JS, Ismail N, Dippenaar A, Ma S, Sherman DR, Warren RM, Kana BD. 2020. Genetic Diversity in Mycobacterium tuberculosis Clinical Isolates and Resulting Outcomes of Tuberculosis Infection and Disease. Annu Rev Genet doi:10.1146/annurev-genet-022820-085940.

50. Homolka S, Niemann S, Russell DG, Rohde KH. 2010. Functional genetic diversity among Mycobacterium tuberculosis complex clinical isolates: delineation of conserved core and lineage-specific transcriptomes during intracellular survival. PLoS Pathog 6:e1000988.

51. Stewart GR, Snewin VA, Walzl G, Hussell T, Tormay P, O’Gaora P, Goyal M, Betts J, Brown IN, Young DB. 2001. Overexpression of heat-shock proteins reduces survival of Mycobacterium tuberculosis in the chronic phase of infection. Nat Med 7:732–7.

52. Wu S, Howard ST, Lakey DL, Kipnis A, Samten B, Safi H, Gruppo V, Wizel B, Shams H, Basaraba RJ, Orme IM, Barnes PF. 2004. The principal sigma factor sigA mediates enhanced growth of Mycobacterium tuberculosis in vivo. Mol Microbiol 51:1551–62.

53. Tollefsen S, Pollock JM, Lea T, Harboe M, Wiker HG. 2003. T-and B-cell epitopes in the secreted Mycobacterium bovis antigen MPB70 in mice. Scand J Immunol 57:151–61.

